# Archaic Adaptive Introgression in Modern Human Reproductive Genes

**DOI:** 10.1101/2024.11.06.622331

**Authors:** Christopher Kendall, Amin Nooranikhojasteh, Esteban Parra, Michael A. Schillaci, Bence Viola

## Abstract

Modern humans and archaic hominins, namely Denisovans and Neanderthals, have been demonstrated to have a long history of admixture. Specifically, some of these admixture events have been adaptive and allowed modern humans to adapt to their new environments outside of Africa. Little research has been done on the impact of archaic introgression on genes associated with reproduction. In this study we report evidence of putative adaptive introgression of 118 genes within modern humans that have been previously associated with reproduction in mice or modern humans. From these genes we identified 11 archaic core haplotypes, three that have been positively selected. Additionally, we found that 327 archaic alleles have genome-wide significance for a variety of traits and 308 of these variants were discovered to be eQTLs regulating 176 genes. We highlight that 81% of the archaic eQTLs overlapping a core haplotype region regulate genes expressed in ovaries, prostate, testes, and vagina compared to other tissues. We also found that several of the putatively adaptively introgressed genes in our results are enriched in developmental and cancer pathways. Further, some of these genes have been associated with embryo development and reproductive-inhibiting phenotypes like preeclampsia. Lastly, we found that archaic alleles overlapping an introgressed segment on chromosome 2 are protective against prostate cancer. Taken together, our results describe how archaic haplotypes, when introduced into a modern human background, may be important in regulating development across the lifespan of an individual.

## Introduction

Modern humans, Denisovans, and Neanderthals share an evolutionary history, with a common ancestor 550,000-765,000 years ago (Meyer et al., 2016). Anatomically modern humans (*Homo sapiens*) evolved in Africa about 300,000 years ago (Hublin et al., 2017) and began the first of many expansions out of Africa by at least 85,000 years ago (Groucutt et al., 2018). Upon dispersal, they encountered and interbred with archaic *Homo*, the Neanderthals and Denisovans, episodically in diverse geographic regions (Green et al., 2010; Reich et al. 2010; Prüfer et al., 2014; Vernot & Akey, 2014; Vernot & Akey, 2015; Kuhlwilm et al., 2016; Browning et al., 2018; Jacobs et al., 2019; Villanea & Schraiber, 2019; Li et al., 2024). As a result of these admixture events in the past, estimates show that on average most non-African populations have just less than 2% of their DNA composed of Neanderthal sequence (Li et al., 2024). Estimates of Denisovan contributions to some human populations tends to be less than 1% on average (Qin & Stoneking, 2015; Sankararaman et al., 2016), save for some Oceanic populations that have been shown to have nearly 5% of their genome shared with Denisovans (Reich et al., 2010; Skov et al., 2018; Jacobs et al, 2019). However, the highest level of Denisovan ancestry recovered to date occurs in the Philippine Ayta (Larena et al., 2019).

The consequences of these admixture events within the context of modern human evolution have been mixed. Some archaic DNA segments have been shown to be adaptively introgressed into modern human populations, where these sequences, conferring some fitness benefit, saw rapid expansion within some groups to high frequencies in relatively short evolutionary timespans.

These include, but are not limited to, several immunity genes (Abi-Rached et al., 2011; Racimo et al., 2015; Dannemann et al., 2016; Vespasiani et al., 2022), keratin genes related to skin and hair phenotypes (Sankararaman et al., 2014; Vernot & Akey, 2014; Dannemann & Kelso, 2017; McArthur et al., 2021), and high-altitude adaptation in Tibetan populations (Huerta-Sánchez et al., 2014). Contrary to this, research over the past decade has shown that introgression deserts exist within the modern human genome, suggestive of removal of archaic segments in the genome (Sankararaman et al., 2014; Sankararaman et al., 2016; Vernot et al., 2016; Skov et al., 2020). Many of these loci were thought to be weakly deleterious and purged via purifying selection in modern human populations (Harris & Nielsen, 2016; Juric et al., 2016). Direct evidence of this reduction in archaic sequences has been documented, where examination of 51 ancient modern human samples with ages between 45,000 and 7,000 years old showed a steady decline of Neanderthal ancestry towards the younger samples, consistent with purifying selection (Fu et al., 2016).

Some research has explored the potential phenotypic expression of this purifying selection. Among these, one compelling discovery was a greater than expected number of archaic loci being removed from the modern human X chromosome relative to the other chromosomes, which may be evidence for male sterility in AMH-archaic hybrids (Sankararaman et al., 2014; Sankararaman et al., 2016). Further evidence comes from research showing that a Neanderthal Y chromosome haplotype has not been documented in modern humans so far, and mutations in this region have been associated with spermatogenesis, also suggesting male sterility of hybrids may have occurred (Mendez et al., 2016). Support for this can be found in Haldane’s Rule, stating that in instances of hybridisation, chromosomal pair fusion difficulties would manifest in a sterility phenotype in the heterogametic sex (Haldane, 1922). However, more recent research has explored reproductive phenotypes, highlighting that archaic populations and modern human groups share some deleterious variants within genes associated with reproduction (Greer et al., 2021). Through modelling, they found that reproductive incompatibilities between modern humans and archaics likely did not occur (Greer et al., 2021). Lastly, research focusing specifically on the impact of loci associated with reproduction derived from Neanderthals into modern humans have found mixed results. For example, Li et al. (2018) documented a missense variant, rs1042838, within the *PGR* locus associated with preterm birth that derived from Neanderthals in some European populations at frequencies as high as 18%. Later analysis showed that a Neanderthal haplotype within the *PGR* gene was linked with a reduction in the number of miscarriages and decreased bleeding during pregnancy, which suggests that within a modern human genetic background, this haplotype may increase fertility (Zeberg et al., 2020).

The research we present here examines the extent of adaptive introgression within genes related to reproduction in modern human populations. While prior analysis has demonstrated both beneficial and deleterious consequences as a result of admixture, to our knowledge, there has been no widespread analysis of the role introgression has played within genes related to reproduction within modern human populations. We speculate that, like the evidence from *PGR* (Zeberg et al., 2020), other archaic-derived loci within genes related to reproduction may show functional benefits in modern human populations, being brought to higher frequencies suggestive of adaptive introgression within specific populations. To explore this, we took advantage of recent high coverage, phased genomic data from the gnomAD 1000 Genomes and Human Genome Diversity Project (1KGP + HGDP) callset containing 76 worldwide populations (Koenig et al., 2024). We examined these populations for evidence of archaic introgression using Neanderthal and Denisovan genomes, including the previously published high coverage Altai Neanderthal (Prüfer et al., 2014), Chagyrskaya Neanderthal (Mafessoni et al., 2020), Vindija Neanderthal (Prüfer et al., 2017), and Altai Denisovan (Reich et al., 2010). We also explore evidence of adaptive introgression within genes related to reproduction in both mice and modern humans listed in Greer et al. (2021).

## Materials and methods

### Identification of archaic segments

We used SPrime (Browning et al., 2018) with the standard settings to identify regions likely of archaic origin within modern human populations. To accurately identify these segments, SPrime uses a scoring algorithm to flag putatively introgressed segments, where it is recommended to filter out segments that score below 150,000 (Browning et al., 2018). Our modern human data came from the recently released phased gnomAD 1KGP + HGDP callset (Koenig et al., 2024), a high-resolution data panel combining the Human Genome Diversity Project (n=51 populations) and 1000 Genomes Project (n=25 populations) mapped to GRCh38 (hg38) coordinates. To filter our data, we followed the SPrime protocol described by Zhou and Browning (2021) by first merging an African reference population, in our case the Yoruba from Ibadan, Nigeria (YRI) population from the 1KGP (n=108 samples), with each of the other populations within our analysis set. To effectively identify putative archaic segments, SPrime requires a non-admixed reference panel to compare the target panel against, which it uses to mask potentially ancestral alleles from being recognised as of archaic ancestry (Browning et al., 2018). We updated the variant IDs for each SNP using the dbSNP database annotation files (Sherry et al., 2001) for known variants so we could match variants in downstream analyses. We kept only biallelic markers after filtering using BCFtools v1.13 (Danecek et al., 2021) and removed any duplicated variants using PLINK2 (Chang et al., 2015).

Our modern human data is mapped to hg38 coordinates, therefore, we used the UCSC LiftOver Linux executable (Hinrichs et al., 2006) to convert our results to the GRCh37 (hg19) genome build, as that matches the published Neanderthal and Denisovan genomes. In case LiftOver (Hinrichs et al., 2006) had mapped any variants incorrectly, we removed variants that had moved from their original chromosomes. We used the Altai Denisovan (Reich et al., 2010), Altai Neanderthal (Prüfer et al., 2014), Vindija Neanderthal (Prüfer et al., 2017), and Chagyrskaya Neanderthal (Mafessoni et al., 2020) as our archaic samples to explore adaptive introgression within human populations. The archaic genomes and their mask files were downloaded from their hosting sites and are provided in the data availability statement. Next, we used map_arch (Zhou & Browning, 2021) to match our results to the archaic samples. This software takes as input the SPrime output, an archaic VCF and associated mask file to list, for each variant in the SPrime output, whether the modern human marker is a match, mismatch, or is not comparable relative to an archaic sample. To generate allele frequencies for the modern human samples, we used BCFtools (Danecek et al., 2021) and subsequently merged these allele frequencies with our initial output using the dplyr package (Wickham et al., 2023) in R v4.1.2 (R Core Team, 2023). Using the output file, we were able to determine the archaic allele and generated the allele frequency of this allele within our population of interest (hereafter archaic allele frequency).

### Dataset of genes associated with reproductive traits

Our research focused on identifying if any archaically-derived segments overlapped genes known to be involved in reproductive traits. To do this we downloaded the list of 1,692 genes found on autosomes which were examined in Greer et al. (2021). These genes potentially impact 11 different biological processes related to reproduction in either mice or modern humans and include: epididymis development or functionality, fertilisation, foetal development, oogenesis, ovary development or functionality, pre- and post-implantation embryo development, sex determination, spermatogenesis, testis development or functionality, and uterus development or functionality (Greer et al., 2021). We focused on autosomal loci because our archaic samples are all female (Reich et al., 2010; Prüfer et al., 2014; Prüfer et al., 2017; Mafessoni et al., 2020), therefore lacking a Y chromosome, and because previous analyses have already identified a dearth of archaic segments on the modern human X chromosome (Sankararaman et al., 2014; Sankararaman et al., 2016; Skov et al., 2020). We aimed to normalise our results by first reporting on variants rather than general segments, and secondly, describing our results using hg19 coordinates to match that of the archaic samples. We used the dplyr package (Wickham et al., 2023) in R (R Core Team, 2023) to extract variants overlapping our genes of interest in our results by matching the variant rsIDs.

### Filtering of archaic segments

We selected segments within our results that have a match/mismatch ratio (the ratio of the number of matches divided by the total number of matches and mismatches found by map_arch) of >50% for the Neanderthals and >40% for the Denisovan (Browning et al., 2018). After removing segments that failed this step, we annotated our VCF files with SnpEff (Cingolani et al., 2012) allowing us to merge gene and consequence data with our population files based on matching the rsIDs. We used BEDTools (Quinlan & Hall, 2010) to intersect the results from each population against every other population to search for potential regional signatures within our results. Additionally, because it is possible that one archaic sample may harbour a signal absent in the other archaic populations, we generated intersection data for every modern human population relative to the four archaic samples in our analysis. To locate regions of contiguous blocks of introgression within our results, we extracted the putatively introgressed segments identified in our analysis from each population that had at least one variant overlapping a gene listed by Greer et al. (2021). Following this, we combined these segments together with populations from the same region according to the gnomAD sample metatable, and reduced them to the minimum number of non-overlapping segments using the GenomicRanges package v3.19 (Lawrence et al., 2013) in R (R Core Team, 2023). We also repeated this analysis to generate a global introgression map regarding our results, where we combined all of the archaic segments regardless of geographic origin in our analysis and reduced them to non-overlapping segments.

### Introgressed segment affinity

Match/mismatch ratios have previously been used to gauge the similarity of genomic segments to one archaic sample compared to the others (Browning et al., 2018; Zhang et al., 2021; Zhou & Browning, 2021). Introgressed segments with 30 or more variants with match/mismatch ratios over 60% to a Neanderthal sample with a concomitant match/mismatch ratio below 40% to the Denisovan are likely of Neanderthal affinity (Browning et al., 2018). Similarly, segments most likely of Denisovan origin will showcase match/mismatch ratios greater than 40% to the Denisovan and less than 30% to the Neanderthals (Browning et al., 2018). To explore this, we assessed which segments are clearly introgressed from one archaic population relative to the others. When a segment passed these donor thresholds with a match/mismatch ratio 5% higher than the other three archaic samples, we suggest the segment is derived from that archaic population. If the match/mismatch ratio is less than 5% higher than the other archaic samples, we consider this segment to be general archaic ancestry. We applied these calculations to our core haplotypes and visualised them using contour plots based on the scripts provided by the SPrime Protocol (Zhou & Browning, 2021) using the kde2d function from MASS (Venables & Ripley, 2002) in R (R Core Team, 2023).

### Identification of high-frequency archaic segments and creation of core haplotypes

Next, we filtered out any variants with archaic allele frequencies below 40% and identified the maximum archaic allele frequency per segment. To generate a series of core haplotypes within our results, we identified variants with allele frequencies of at least 40% that directly matched the archaic allele and were found within one of the genes listed in Greer et al. (2021) described as being associated with reproductive phenotypes. We focused this analysis on populations from the 1KGP because of the small sample sizes of the HGDP, with the exception of the Melanesian and Papuan population samples of the HGDP, so we could investigate potential Denisovan contributions. Once we located the maximum archaic allele in each segment, we kept variants that are no more than 5% below the maximum archaic allele frequency directly upstream and downstream from the maximal variant, discarding the rest of the segment for this specific analysis. The remaining windows around the maximum archaic allele are what we consider to be core haplotypes within our analysis.

### Detection of positive selection

Additionally, we wanted to explore for evidence of positive selection in our results, so we used RAiSD (Alachiotis & Pavlidis, 2018) with standard input parameters to scan for signatures of positive selection within our population-specific segments that contain a core haplotype. Variants determined to be under positive selection are those with scores greater than 99.5% of the other variants within their chromosome, a recommended cut-off described in the RAiSD documentation (Alachiotis & Pavlidis, 2018).

### Haplotype networks and ancestral recombination graphs

We generated a haplotype network for all of our modern human populations and the archaic samples for the *CSNK1A1-ENSG00000230551* core haplotype, as this core displayed evidence of positive selection (see Matrials and Methods). Because haplotypes require phased data, and the archaic samples are not phased, we took advantage of the fact that our Neanderthal and Denisovan samples have very long runs of homozygosity (Sánchez-Quinto & Lalueza-Fox, 2015; Prüfer et al., 2017; Mafessoni et al., 2020; Villanea et al., 2021). Therefore, removing heterozygous sites from these samples will artificially phase the genomes because both alleles will be the same. Further, prior analyses have made use of a similar approach and were able to generate effective networks showing haplotype similarities between Denisovans and Tibetan populations (Huerta-Sánchez et al., 2014). After filtering out heterozygous sites using BCFtools (Danecek et al., 2021) on chromosome 5, we removed 22,976 sites (0.0001926% of the total sites on chromosome 5) from the Altai Neanderthal, 17,938 sites (0.0001504% of the total sites on chromosome 5) from the Chagyrskaya Neanderthal, 26,736 sites (0.0002242% of the total sites on chromosome 5) from the Denisovan, and 20,533 sites (0.0001721% of the total sites on chromosome 5) from the Vindija Neanderthal. As a consequence, from the *CSNK1A1-ENSG00000230551* core haplotype we kept all sites in the Altai Neanderthal core and removed one site (0.00001681% of the total variants) from the Chagyrskaya Neanderthal core, two sites (0.00003363% of the total variants) from the Denisovan core, and 16 sites (0.0002690% of the total variants) from the Vindija Neanderthal core. In total, we were left with 1,009 variants over a 98,609 base pair region for analysis.

We generated a FASTA format file for *CSNK1A1-ENSG00000230551* from the merged VCF using PGDSpider (Lischer & Excoffier, 2012) followed by using MUSCLE v3.8.31 (Edgar, 2004) to align the file. We converted the aligned FASTA file to NEXUS format (Maddison et al., 1997) using MEGA11 (Tamura et al., 2021), and then imported the file into DnaSP6 (Rozas et al., 2017), where we assigned population sequence sets and generated a haplotype file. PopART v1.7 (Leigh & Bryant, 2015) was used to build the haplotype network using the median-joining function method. We combined populations according to their region/superpopulation (i.e. AFR/African, AMR/American) in our haplotype file, traits file, and haplotype network because of the large number of subpopulations we included in our analysis. In total, *CSNK1A1-ENSG00000230551* has 965 haplotypes. To make our haplotype network legible, we included only the top 50 most frequent haplotypes, and their ties, along with the haplotypes associated with the archaic samples. This left us with 54 haplotypes to plot.

We used Relate v1.2.1 to generate ancestral recombination graphs (Speidel et al., 2019). To do this, we first generated a new VCF from the gnomAD 1KGP + HGDP phased callset (Koenig et al., 2024) with all of the populations present and converted the VCF to the proper format using the functions native in Relate, including the RelateFileFormats --mode ConvertFromVcf and PrepareInputFiles.sh commands (Speidel et al., 2019). Once filtering was completed, we were left with 2,339 variants for further analysis within the *CSNK1A1-ENSG00000230551* core haplotype region. To run Relate, we used the recommended settings of a mutation rate of 1.25e^-8^ and an effective population size of 30,000. The trees we selected for further review included derived branches originating at least 1,000,000 years ago, with subsequent long, non-recombining branches, followed by rapid expansion very recently. We used Relate to extract subpopulation trees (RelateExtract --mode SubTreesForSubpopulation) to investigate the genealogies in more detail on a global scale for variants with an archaic-looking tree that also were labelled as matches in map_arch (Zhou & Browning, 2021). Following this, we plotted mutational trees (TreeViewMutation.sh) using the built in Relate functions with the YRI as our outgroup and representative of the rest of the sub-Saharan African populations to generate legible trees.

### Identification of archaic variants showing genome-wide significant associations with common traits and diseases

We were interested in determining if any of the archaic variants in our results show genome-wide significant associations (p=5×10^-8^ or less) with complex traits and diseases. We compared the variants in our results against the IEU OpenGWAS project database (Hemani et al., 2018; Elsworth et al., 2020). For any matches that were found at genome-wide significance, we downloaded the associated genome-wide association study (GWAS) summary statistics and compared these to our archaic alleles. If the effect allele did not match the archaic allele, we converted the direction of effect (beta) by flipping the sign and then updated the non-effect allele.

### Gene expression and regulation

We explored whether archaic variants showing genome-wide significant associations in our results were involved in the regulation of gene expression. Using FUMA GWAS’s SNP2GENE function (Watanabe et al., 2017; Watanabe et al., 2019) we input the variants compiled in our OpenGWAS (Hemani et al., 2018; Elsworth et al., 2020) results (n=327) and then downloaded the expression quantitative trait loci (eQTL) table from the output. This process was repeated again just for significant variants that overlapped our core haplotypes (n=114).

### Variant annotations and gene ontology

Variant annotation was done using the GRCh37 search in SNPnexus (Chelala et al., 2009; Oscanoa et al., 2020) to understand the functional consequences of archaic variants identified in our data (n=880). Further, we used BioMart’s GRCh37 Release 112 (Harrison et al., 2024) to access gene ontology (GO) GO Consortium information (Ashburner et al., 2000; Gene Ontology Consortium et al., 2023) for the genes with high-frequency archaic segments identified in our results (n=118).

### Enrichment analysis

We ran several analyses to determine if any of the genes found within our results were enriched in any pathways. We used ShinyGO (Ge et al., 2020) to test for significantly enriched KEGG pathways (Kanehisa, et al., 2023) for the genes which contain genome-wide significant markers (n=47). We supplemented this with results from Enrichr (Chen et al., 2013; Kuleshov et al., 2016; Xie et al., 2021), which scans the Reactome 2022 (Milacic et al., 2024) and KEGG 2021 human (Kanehisa, et al., 2023) pathways, and the GO biological processes and GO molecular functions (Ashburner et al., 2000; Gene Ontology Consortium et al., 2023) databases for any other significant pathways not outlined in our first analysis. We repeated these analyses just for the genes overlapping core haplotype segments with genome-wide significant markers (n=10). Lastly, we also performed enrichment analysis on the set of genes regulated by archaic eQTLs identified by FUMA GWAS (Watanabe et al., 2017; Watanabe et al., 2019) for all genes (n=234) and just those regulated within the core haplotypes (n=48).

## Results

### Segments with high-frequency archaic variants

We found 47 archaic segments collectively in global modern human populations overlapping genes associated with reproduction. These segments represent 37.88Mb (million base pairs/megabases) of sequence data with at least one variant in each segment surpassing a frequency of 40% (Supplemental Table 1). Regarding regional-specific segments suggestive of adaptive introgression, we found 26 segments in American, 17 in East Asian, 6 in European, 1 in Middle Eastern, and 6 in Oceanic populations, respectively (Supplemental Table 1). We extracted the 47 global segments from each of our populations and grouped them by region to visualise the admixture relationships between modern humans and the archaic samples within these specific regions. Even with our specific focus on only genes associated with reproduction, we were able to recreate many of the previously established introgression patterns in modern human groups from archaic populations. These include the Chagyrskaya and Vindija Neanderthals sharing more alleles with the introgressing Neanderthal sequence in modern humans (Prüfer et al., 2017; Mafessoni et al., 2020) and Oceanic populations having a higher proportion of Denisovan ancestry than other global populations (Reich et al., 2010; Skov et al., 2018; Jacobs et al, 2019). As expected, African populations have considerably less introgressed segments relative to non-Africans (Li et al., 2024). These relationships are shown in Figure 1. The list of population specific segment containing archaic alleles with frequencies of at least 40% can been in Supplemental Table 2.

**Figure 1.**
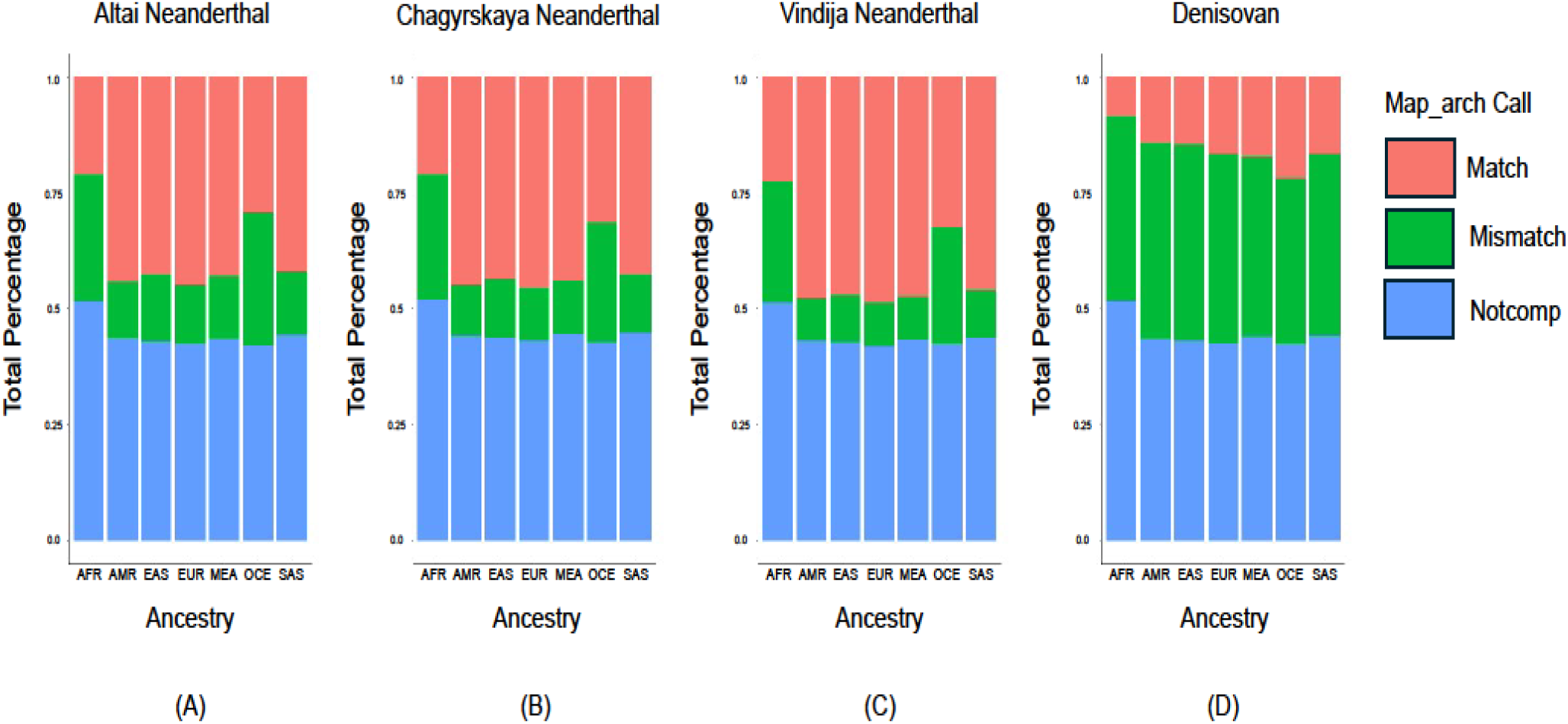
Variants recovered in adaptively introgressed reproductive genes. Putative archaic variants recovered from segments overlapping segments from adaptively introgressed reproduction-associated genes. The colours denote map_arch labelling (Zhou & Browning, 2021) and are representing the percentage of total recovered variants for the (A) Altai Neanderthal, (B) Chagyrskaya Neanderthal, (C) Vindija Neanderthal, and (D) Denisovan archaic samples. The population codes refer to: African (AFR), American (AMR), East Asian (EAS), European (EUR), Middle Eastern (EAS), Oceanic (OCE), and South Asian (SAS). The plots were generated using the ggplot2 package (Wickham, 2016) package in R (R Core Team, 2023). A high resolution version of this image is available at https://doi.org/10.5281/zenodo.14047487.

### Core haplotypes

We generated core haplotypes based on archaic variants that had allele frequencies ≥ 40% in the 1KGP or Melanesian/Papuan HGDP samples, directly matched the archaic allele, were the maximum archaic allele frequency within their segment, and were found within one of the 1,692 genes described in Greer et al. (2021). After filtering for this criteria, we were able to generate 11 core haplotypes that overlap 15 genes (Supplemental Table 3). Three of these segments have evidence of positive selection. The *CSNK1A1-ENSG00000230551* region (chr5:148869994-149203782) in the Han Chinese South (CHS) population shows positive selection within the main core haplotype. Another two segments, chr13:28962942-28997886 in the Peruvian in Lima Peru (PEL), overlapping the gene *FLT1*, and chr20:50563299-51218753 in the Papuans, overlapping the gene *SALL4*, show evidence of positive selection within the larger introgressed segment generally, but outside of the core haplotype (Supplemental Table 3). *CSNK1A1-ENSG00000230551* and *FLT1* were found to have the most similarity with the Vindija Neanderthal while *SALL4* shows the highest similarity with the Denisovan (Supplemental Table 3). We created a haplotype network and generated ancestral recombination graphs for *CSNK1A1-ENSG00000230551* due to the discovery of the positive selection within its core segment. In total, we identified 965 haplotypes within the *CSNK1A1-ENSG00000230551* core region. The top 50 haplotypes and their ties for *CSNK1A1-ENSG00000230551*, along with the haplotypes of the four archaic samples, are plotted in Figure 2 while ancestral recombination graphs for this region are shown in Figure 3. The regions where RAiSD (Alachiotis & Pavlidis, 2018) detected positive selection are presented in Supplemental Table 3 when selection exists. Lastly, we generated contour plots (Zhou & Browning, 2021) for the *CSNK1A1-ENSG00000230551* core haplotype displaying the relationships to each of the archaic samples, which is shown in Figure 4. To add onto this, we list in Supplemental Table 3 the most probable archaic donor for each of our core haplotypes.

**Figure 2.**
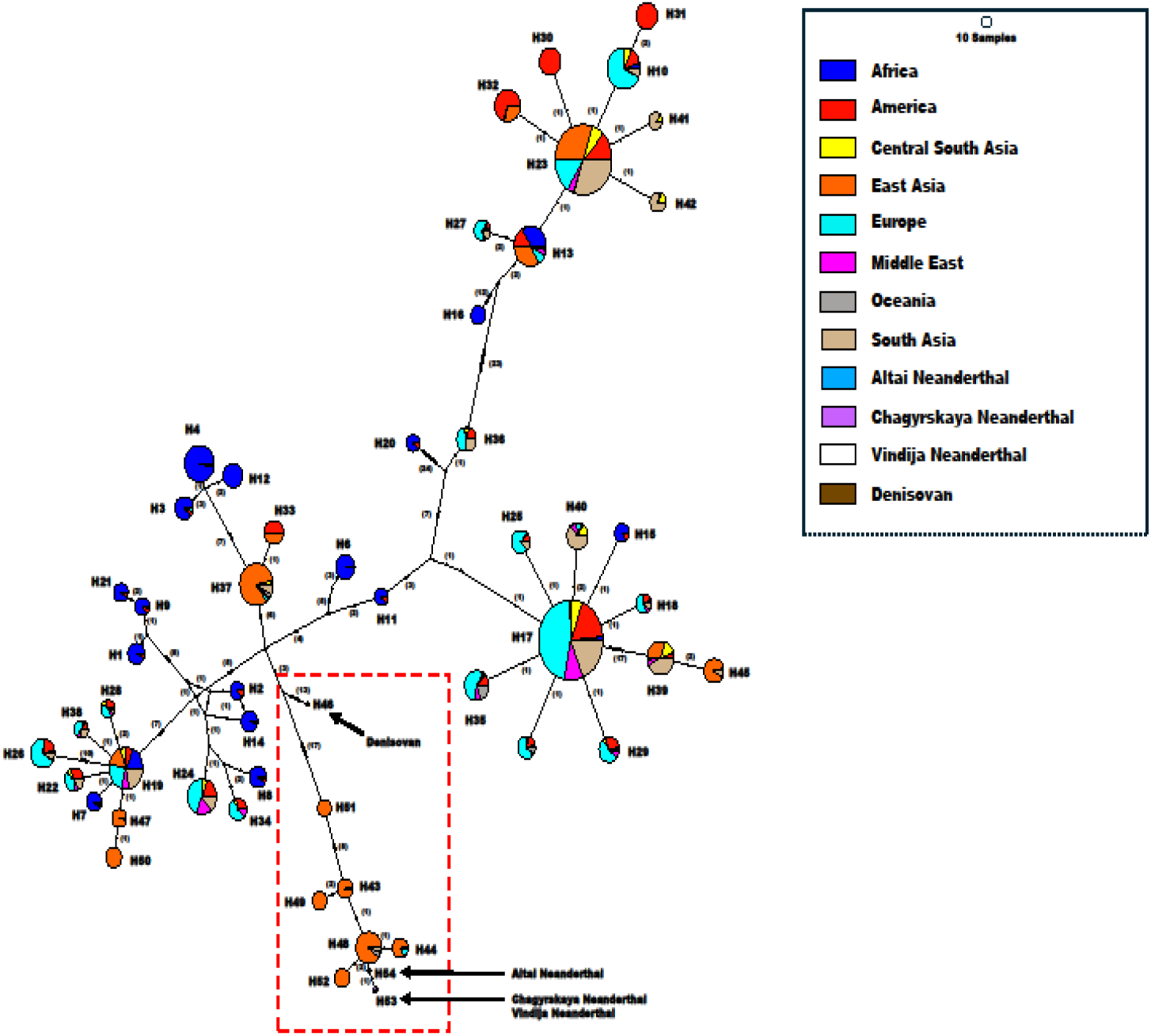
Haplotype network for *CSNK1A1-ENSG00000230551*. The haplotype network plot was generated using PopArt v1.7 (Leigh & Bryant, 2015) for *CSNK1A1-ENSG00000230551* our core haplotype with evidence of positive selection within its core region. The red box highlights the location of the archaic haplotypes and the number of mutations along each edge between nodes is shown in brackets. A high resolution version of this image is available at https://doi.org/10.5281/zenodo.14047487.

**Figure 3.**
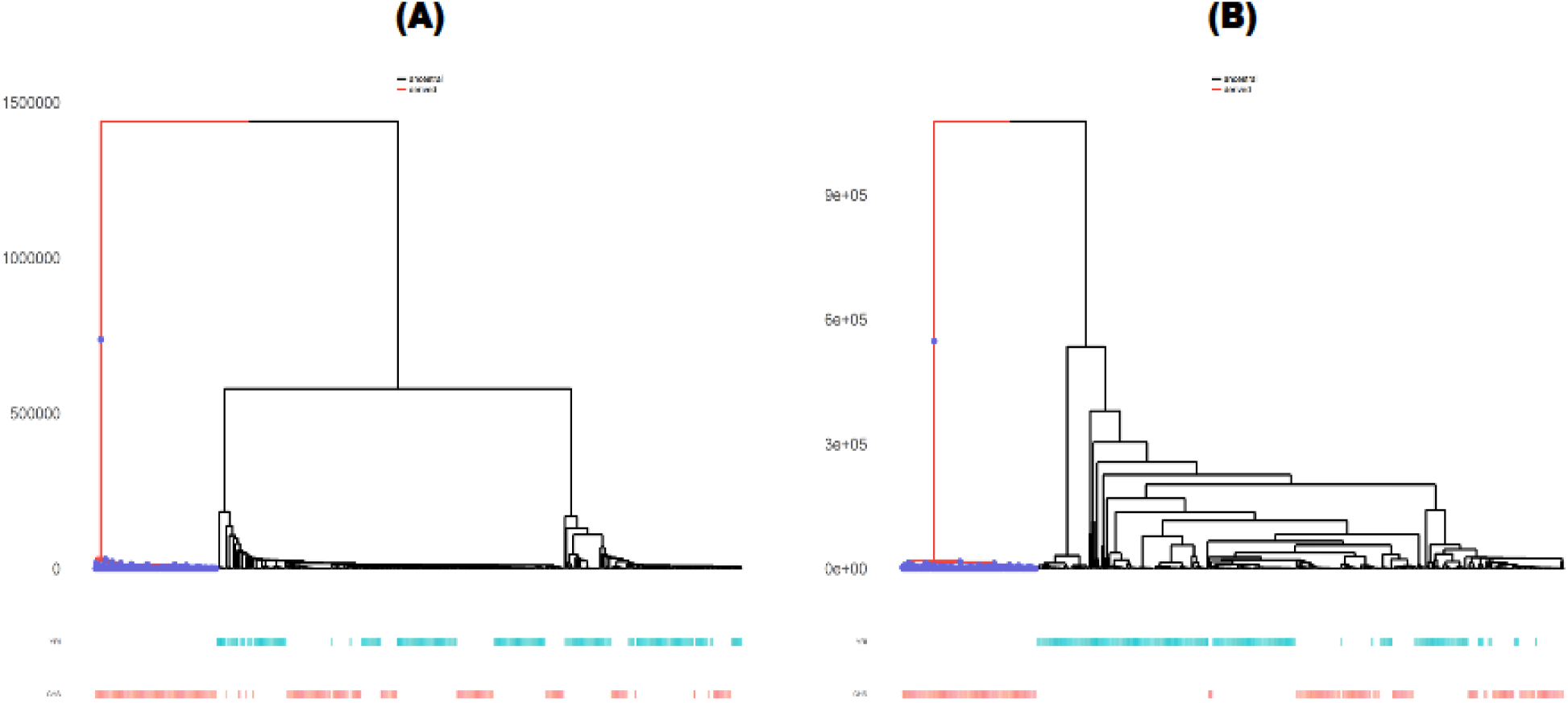
Ancestral recombination graphs of *CSNK1A1-ENSG00000230551*. Ancestral recombination graphs for variants within the *CSNK1A1-ENSG00000230551* core haplotype that are matches to the archaic allele and show patterns associated with archaic ancestry. These patterns include origins at least 1,000,000 years ago and long, non-recombining branches with recent expansion within modern human populations associated with derived mutations. (A) rs76352267 (chr5:148869994) and (B) rs12055320 (chr5:148876726) inside of the CHS population, which show archaic-like branching events. The analysis and graphs shown here were generated using Relate v1.2.1 (Speidel et al., 2019). A high resolution version of this image is available at https://doi.org/10.5281/zenodo.14047487.

**Figure 4.**
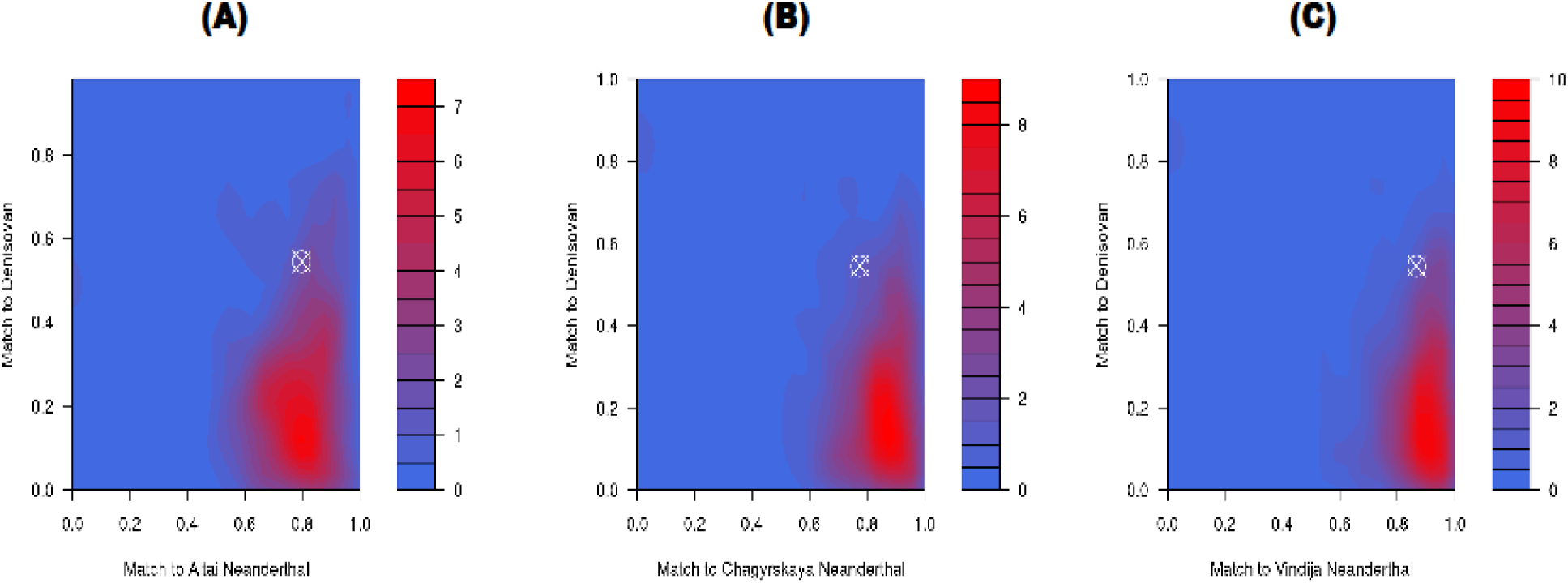
Contour plots for *CSNK1A1-ENSG00000230551* core haplotype. The contour plots based on the match/mismatch ratios of authentic introgressed segments. The location of the segment containing the gene is identified by the white crosshair. The density plot is based on the number of segments genome-wide from both the Neanderthal sample (x-axis) and Denisovan (y-axis) overlapping each match/mismatch ratio, with the archaic donor population being the archaic sample with the highest match/mismatch ratio. (A) Altai Neanderthal vs. Denisovan, (B) Chagyrskaya Neanderthal vs. Denisovan, and (C) the Vindija Neanderthal vs. Denisovan, with the Vindija Neanderthal being the archaic sample with the highest match rate. However, it should be noted that the Denisovan also shares very high levels of ancestry here as well (over 40%), making it difficult to determine if this is truly of Neanderthal affinity despite the high similarity to the Vindija sample, or just of archaic origin generally. The plots were based upon the scripts provided by Zhou and Browning (2021) using the kde2d function from the MASS library (Venables & Ripley, 2002) in R (R Core Team, 2023). A high resolution version of this image is available at https://doi.org/10.5281/zenodo.14047487.

### Association of archaic variants with GWAS traits and gene expression

Our identified segments (Supplemental Table 1) overlap 880 variants within 118 genes and intergenic segments (Supplemental Table 2). Of these variants, 327 of them, located within 47 genes, are found at genome-wide significant thresholds. These results are listed in Supplemental Table 4. From this larger list of variants, 114 of them were found to overlap 10 of our core haplotype segments. Next, we identified variant associations by running SNPnexus for functional consequences, which we provide in Supplemental Table 5. Lastly, we wanted to see if any of our genome-wide significant variants (Supplemental Table 4) were associated with gene expression levels (i.e. were eQTLs) and used the SNP2GENE function within FUMA GWAS (Watanabe et al., 2017; Watanabe et al., 2019) to analyse these variants. In total, 308 variants were discovered to be eQTLs regulating 176 genes, while 113 variants regulating 44 genes were found within our core haplotype regions. Supplemental Table 6 lists the full list of eQTLs while Supplemental Table 7 lists the eQTLs from the core haplotypes.

### Gene enrichment ontologies and pathways

For all genes in our dataset, we used BioMart (Harrison et al., 2024) to explore gene ontology associations generally, which are included in Supplemental Table 8. Expanding on this analysis we further examined our genes to see if they were significantly enriched in any pathways or ontologies. We did this in four separate runs, by first including all genes from Supplemental Table 2 with genome-wide significant variants (set 1), core haplotype genes with genome-wide significant variants (set 2), all genes that are regulated by archaic eQTLs (set 3), and genes that are regulated by archaic eQTLs found within a core haplotype region (set 4). We used ShinyGO (Ge et al., 2020) and Enrichr (Chen et al., 2013; Kuleshov et al., 2016; Xie et al., 2021) to examine each of the four datasets above.

Within set 1, we found no significant ontologies, however, 9 pathways were significantly enriched in our ShinyGO (Ge et al., 2020) results, 5 were significantly enriched in our Reactome 2022 results (Milacic et al., 2024), and two significantly enriched in the KEGG 2021 human (Kanehisa, et al., 2023) pathways identified by Enrichr (Chen et al., 2013; Kuleshov et al., 2016; Xie et al., 2021). The ShinyGO results for set 1 can be seen in Supplemental Table 9, while the Enrichr results can be seen in Supplemental Table 10. For set 2, ShinyGO (Ge et al., 2020) and Enrichr (Chen et al., 2013; Kuleshov et al., 2016; Xie et al., 2021) were both unable to identify any significant pathways. However, Enrichr (Chen et al., 2013; Kuleshov et al., 2016; Xie et al., 2021) located 58 significant GO biological processes and 10 significant GO molecular functions. The results for set 2 are listed in Supplemental Table 11.

In set 3, there were no significant ontologies, or any significant pathways found according to Enrichr (Chen et al., 2013; Kuleshov et al., 2016; Xie et al., 2021). Our ShinyGO (Ge et al., 2020) analysis for set 3, alternatively, was able to identify three significant pathways, which included carbon metabolism, central carbon metabolism in cancer, and DNA replication. These results can be viewed in Supplemental Table 12. Lastly, for set 4, ShinyGO (Ge et al., 2020) and Enrichr (Chen et al., 2013; Kuleshov et al., 2016; Xie et al., 2021) both indicated no significant pathways for these genes. Enrichr (Chen et al., 2013; Kuleshov et al., 2016; Xie et al., 2021) also did not find any significant GO biological processes (Ashburner et al., 2000; Gene Ontology Consortium et al., 2023), but did highlight four significant GO molecular processes, which can be seen in Supplemental Table 13.

## Discussion

### Core haplotypes

Our analysis identified 11 core haplotypes, three of which have evidence of positive selection within their introgressed segment (Supplemental Table 3). In particular, one of these haplotypes, *CSNK1A1-ENSG00000230551*, had evidence of positive selection within the core region. *CSNK1A1* was described in Greer and colleagues (2021) as having an infertility phenotype in mice. Additional roles for *CSNK1A1* include embryogenesis (Elyada et al., 2011; Ma et al., 2023), stem cell expansion, certain cancers (Schneider et al., 2014), endometriosis (Zhou et al., 2021), and infertility due to oocyte loss (Zhang et al., 2022). The haplotype network for this region shows that the three Neanderthals are very similar to a large cluster of East Asian haplotypes (Figure 2). Interestingly, one sample from the Chinese Dai in Xishuangbanna, China (CDX) shares the Chagyrskaya and Vindija Neanderthal haplotype (Figure 2), which is also only one mutational step away from the Altai Neanderthal haplotype. Our ancestral recombination graphs (Figure 3) for this region highlighted two variants with very clear archaic signatures. Both chr5:148876726 and chr5:148869994 have long, non-recombining, derived branches originating approximately 1,000,000 years ago, followed by rapid expansion fairly recently within the CHS, a pattern that is absent in the YRI. Analysis of the maximum archaic allele within the *CSNK1A1-ENSG00000230551* core haplotype also agrees with this assessment as all high frequency variants are isolated within Asia or Oceania in this region (Figure 5A).

**Figure 5.**
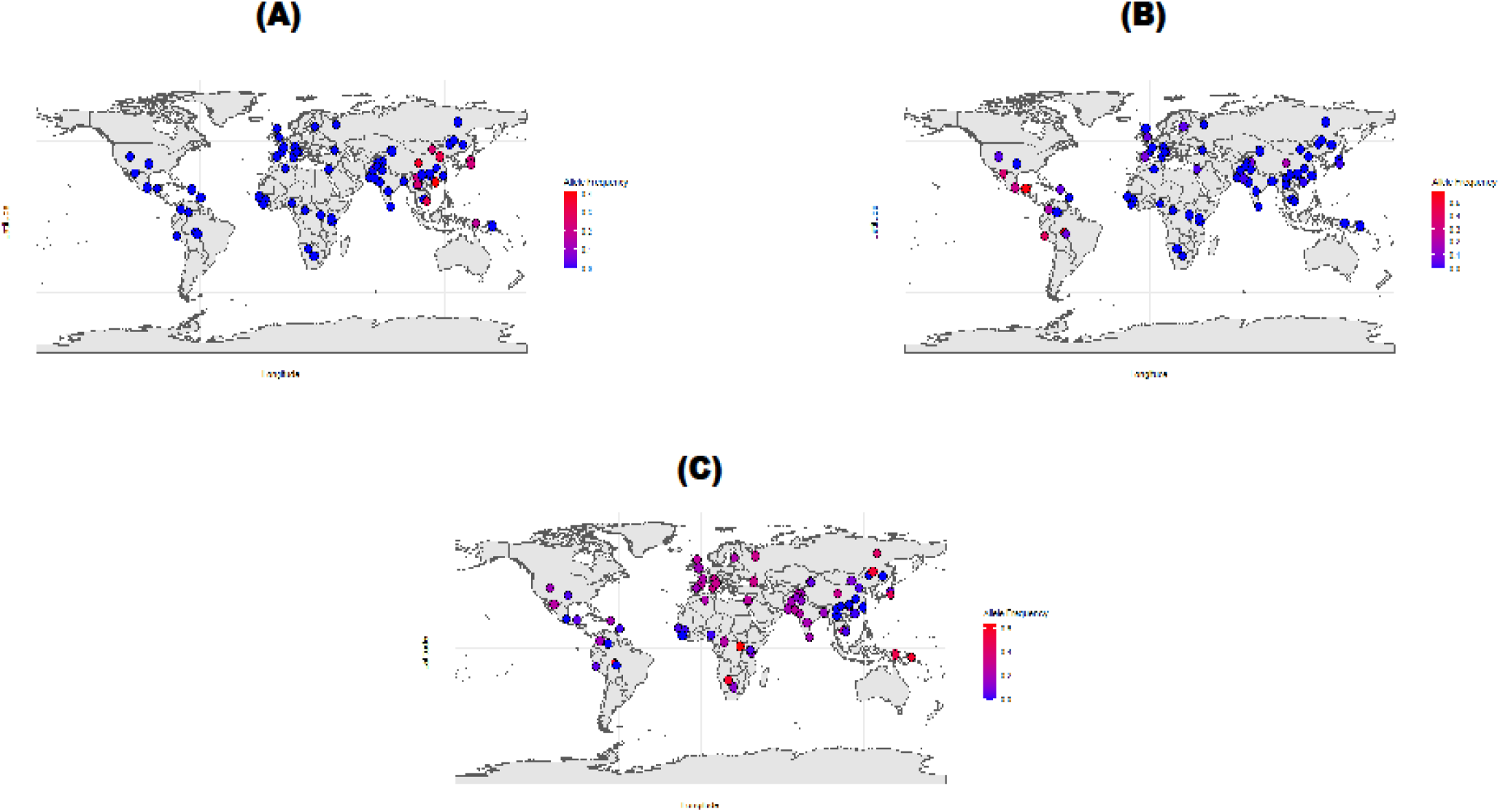
Frequency maps of core haplotypes with evidence of positive selection. Frequency distribution maps generated by extracting the maximum archaic allele frequency from each core haplotype that had evidence of positive selection generally within its putatively derived segment. (A) *CSNK1A1-ENSG00000230551* has high frequency alleles almost exclusively in Asian and Oceania while (B) *FLT1* is almost exclusively in the Americas at high frequency. (C) *SALL4* is seen nearly globally but the highest frequencies are located along the Pacific coast of Asia. The plots were generated with the rnaturalearth (Massicotte & South, 2024), sf (Pebesma, 2018; Pebesma & Bivand, 2023), and ggplot2 (Wickham, 2016) packages in R (R Core Team, 2023). A high resolution version of this image is available at https://doi.org/10.5281/zenodo.14047487.

*FLT1* is a core haplotype that has positive selection generally within the larger introgressing segment (Supplemental Table 3). This gene was included in Greer et al. (2021) due to its association with post-implantation embryo development. Prior research has established a causal marker near *FLT1* linked with preeclampsia (McGinnis et al., 2017; Ashar-Patel et al., 2017; Kikas et al., 2020) and the gene has been speculated as able to negatively regulate angiogenesis in mice, ultimately leading to embryonic death if divergent genotypes are found (Hiratsuka et al., 1998). Interestingly, Arthur and colleagues (2018) suggest that positive selection within the *FLT1* gene in their preeclampsia study may be a result of antagonistic coevolution that drove benefits for the offspring and may have changed genes expressed in the mother ultimately leading to signals of positive selection in both gene sets. Additional phenotypes for the *FLT1* gene include pregnancy loss, low birth weights, and placental inflammation (Muehlenbachs et al., 2008). The geographic distribution of the high frequency alleles is seen most strongly in the Americas within this haplotype, with some slightly elevated frequencies seen within Asia and Europe (Figure 5B). This discovery is consistent with previous research that described signatures of positive selection in the Mexican Ancestry in Los Angeles CA USA (MXL) and Peruvian in Lima Peru (PEL) populations within the *FLT1* gene as a result of archaic admixture (Witt et al., 2023), which is interesting considering *FLTI* was identified as being a core haplotype in the PEL within our results (Supplemental Table 3).

*SALL4* is another core haplotype with positive selection outside of the core (Supplemental Table 3). This gene has been associated with post-implantation embryo development in mice (Greer et al., 2021). In addition to this, mouse models have highlighted roles for *SALL4* in oogenesis, preimplantation embryo development (Xu et al., 2017), along with expression in spermatogonia (Chan et al., 2017), indicating some connection to gamete formation. The maximum archaic alleles within this core haplotype segment are seen almost globally, with the strongest signatures in Oceania, Japan, and Russia (Figure 5C). There are moderate frequencies near the Himalayas and some parts of South Asia that extends into the majority of Europe. Notably, almost all of the East Asian populations and many populations from the Americas have very low variant frequencies in this region.

Unfortunately, there were no genome-wide significant markers overlapping any of these core haplotype segments (Supplemental Table 4) making it difficult to understand the impact archaic introgression may have had on any phenotypes of these genes. There was only one archaic SNP from any of the core haplotypes with evidence of positive selection, rs56314249 overlapping *FLT1*, that was found to be a coding variant. However, the effect was synonymous and generated no amino acid change (Supplemental Table 5), adding another layer of difficulty in interpreting exactly the role of these archaic variants and why their maximum archaic alleles were driven to such high frequencies within our populations. This is especially true considering only one of these core haplotypes comes from the HGDP, which often have small sample sizes. It is difficult to make any inferences regarding these introgressed segments and their direct impact to modern humans, nonetheless, it is noteworthy that both *CSNK1A1* and *FLT1* have been previously identified as containing segments introgressed from archaic populations in several studies (Enard & Petrov, 2018; Silvert et al., 2019; Koller et al., 2022; Gao et al., 2023; Rong et al., 2023; Wei et al., 2023; Witt et al., 2023), confirming our results. The *SALL4* core haplotype region has been previously described through identifying archaic SNPs in the UK Biobank cohort (Dannemann & Kelso, 2017) and additionally in a later introgression map based on Icelandic genomes (Skov et al., 2020). While it is clear that these segments are of archaic origin, and these genes have been tied to both male and female phenotypes associated with reproduction, more research is necessary going forward to properly elucidate the roles the archaic alleles have within this context.

### Adaptively introgressed genes and prostate cancer risk

We identified 15 variants in four genes that have significant associations with prostate cancer risk (Supplemental Table 4) within the chr2:173199359-173589239 segment isolated within the Tujia population (Supplemental Table 2). These genes, *ENSG00000225205*, *ENSG00000226963*, *ITGA6*, and *ITGA6-AS1*, have archaic alleles that reduce the risk of prostate cancer (Supplemental Table 4), reported in large GWAS cohorts from Asia and Europe (Neale lab, 2018; Schumacher et al., 2018; Sakaue et al., 2021; Sato et al., 2023). Previous research has shown that one variant in our analysis, rs12621278, was also found to be protective for prostate cancer onset in Asian populations (Dannemann, 2021). Our enrichment analyses also show that several of the genes with high-frequency archaic segments are important in cancer pathways, where ShinyGO highlighted four significantly enriched cancer pathways within our set 1 genes (Supplemental Table 9) and one significantly enriched pathway, central carbon metabolism in cancer, within set 3 (Supplemental Table 12). Further evidence for our genes being associated with prostate cancer comes from set 2, where one of our core haplotypes, *PPP3R1*, identified in the CDX population overlapping the chr2:68173949-68503920 region, has been shown to be associated with pathways related to prostate cancer previously (Yuan et al., 2015) and is also enriched in several GO biological processes and molecular functions (Ashburner et al., 2000; Gene Ontology Consortium et al., 2023) such as calcium ion binding, calcium- and calcineurin-mediated signaling (Supplemental Table 11). Calcium ions and calcineurin have both been shown to have roles regarding prostate cancer development and growth (Manda et al., 2015; Ardura et al., 2020). Lastly, some genes regulated by core haplotype eQTLs are significantly enriched in palmitoyltransferase activity, which previous research implicated in cancer initiation and modulation (Chen et al., 2024).

### Enrichment for development-related ontologies and pathways

Our results from four genes in set 1 show ties with the Notch signaling pathway (Tables C9 & C10). The Notch signaling pathway was described before as being important for controlling proper foetal development of several vital organs (Reichrath & Reichrath, 2020; Anusewicz et al., 2021). Some of these genes have reproductive associations (Baldarelli et al., 2024). For example, within mouse models, mutant forms of *JAG1* were associated with sterility and premature death (Roderick & Hawes, 1974) and *KAT2B*, a hormone receptor, were differentially expressed in polycystic ovary syndrome (PCOS) oocytes (Liu et al., 2016). Additionally, our core haplotype segments (set 2) identified 68 GO biological processes (Ashburner et al., 2000; Gene Ontology Consortium et al., 2023), 18 of which are related to cardiac, muscle, or skeletal biological processes (Supplemental Table 11). One gene, *TTN*, has displayed various skeletal and cardiac anomalies using mouse models (Baldarelli et al., 2024). This includes an embryonic shrunken head phenotype associated with an inability to generate proper circulation eventually leading to death of the foetus (May et al., 2004) and increased left ventricular diastolic stiffness causing a reduction in activity tolerance (Granzier et al., 2014). In modern humans, disease models show that *TTN* is tied to dilated cardiomyopathy and two forms of muscular dystrophy (Baldarelli et al., 2024). Our GWAS results for *TTN* also support some of the associations addressed here as 21 SNPs overlapping this gene have an archaic allele associated with lower pulse rates (Mitchell et al., 2019) (Supplemental Table 4), which may signal weakening heart abilities. Also for set 2, significant enrichment for calcium ion binding (GO:0005509) was identified (Supplemental Table 11). Calcium is important in mammalian reproduction, having roles in sperm development, embryogenesis, and healthy development of the foetus (Stewart & Davis, 2019). In our set 4 genes, we have significant GO molecular functions related to palmitoyltransferase activity, functions related to neuritogenesis and signaling in early development (Zhang et al., 2017). Taken together, three of the gene sets are significantly enriched in ontologies and pathways that are associated with healthy development of the foetus across several important organs. Additional genes within these sets linked to reproduction suggest that these gene pathways may be important in modulating proper development and function of several important processes across an organism’s lifespan.

### Archaic eQTLs

Approximately 94% of the archaic variants showing genome-wide significant associations with complex traits and diseases (308 out of 327) are also eQTLs (Supplemental Table 6). Over 74% of these eQTLs modulate gene expression in ovaries, prostate, testes, uterus, and vagina. When filtering for eQTLs overlapping with core haplotype segments, this figure is increased to 81% of the eQTL set, save the uterus where no core haplotype eQTLs regulated gene expression in this tissue (Supplemental Table 7). When compared to other important tissues from several major organs combined (brain, liver, lung, spleen, and stomach), only ∼63% of the eQTLs are reported for these tissues. These results suggest that our archaic variants are important in the regulation of gene expression in reproductively-associated tissues. In light of our results above finding enriched pathways associated with cardiac and skeletal development, we also explored expression levels within the heart and musculoskeletal systems. Here, we find approximately the same numbers of eQTLs as we did in the ovaries, prostate, testes, uterus, and vagina, where 73% of the eQTLs are reported for these tissues. In the core haplotypes, the number of eQTLs expressed in heart and musculoskeletal dropped to 69.9% (Supplemental Table 7). Therefore, archaic eQTLs in our analysis, particularly in the core haplotypes, appear to be important in helping regulate genes expressed in the ovaries, prostate, testes, uterus, and vagina with a higher number of eQTLs regulating gene expression in these tissues compared to others.

## Limitations

Our research is not without limitations. Several of the HGDP target populations we analysed here have sample sizes below 15, a recommended minimum number of included individuals for SPrime to maintain high levels of accuracy in identifying introgressed segments (Browning et al., 2018). However, due to sampling bias present in many genomic studies (Hünemeier, 2024), research must rely on populations with limited sample sizes at this time if they want to explore patterns in populations outside of Eurasia. A possible consequence of this is that some of our segments listed in our results may represent false positives, however, the likelihood of this remains small as these segments did pass our filtering parameters to be considered an authentic segment (see Materials and Methods), and further, many of these segments are also shared with populations with adequate sample sizes (i.e. those from the 1KGP) suggesting they are accurately attributed as introgressed. A second consequence of this is elevated allele frequencies in the HGDP, which may not be found in populations with more than 15 samples. We acknowledge that using strictly allele frequency as a measure of adaptive introgression may lead to artificially elevated detection of adaptively introgressed segments in our results. Therefore, we suggest caution in interpreting some of the results from the HGDP due to their low sample sizes and concomitant elevated allele frequencies unless they are paired with higher-than-expected archaic allele frequencies in other populations with adequate sample sizes. Another possible drawback is our use of the SPrime software, which has been shown to mask some instances of admixture if the segments are too similar to that of the human outgroup (Chen et al., 2020). It is possible that using other software that is reference-population-free to repeat our study, such as IBDMix (Chen et al., 2020), may uncover even more interesting results not currently presented in our results. Lastly, we also used GWAS summary statistics to infer the direction of effect of the archaic variants. The interpretation of some of these effects is not always straightforward as the phenotypes reported in some studies may not be clearly defined. Additionally, although a number of archaic alleles display genome-wide significant effects for a wide range of traits, this does not necessarily prove they are the causal variants, and the associations may be simply due to the archaic allele being in linkage disequilibrium with the actual causal variants. Since identifying causal variants was not the aim of this project, future research should explore fine-mapping and functional validation to uncover if the variants we present here are causal in these trait effects or a byproduct of linkage disequilibrium.

## Conclusion

Our study identified 47 independent segments within modern humans harbouring high-frequency archaic variants from Neanderthals and Denisovans, suggesting adaptive introgression. These segments are located within genes that have been previously described as being related to reproduction and fertility in either humans or mice (Greer et al., 2021). In total, these segments overlap 880 variants within 118 genes, where the variants have archaic allele frequencies ≥ 40%. Further filtering of these segments revealed 11 core haplotypes that have at least one variant with an archaic allele frequency ≥ 40%, the variant directly matched the archaic allele, the variant was the maximum archaic allele frequency within its introgressed segment, and the variant was found within one of the 1,692 genes described in Greer et al. (2021). Three of these core haplotypes displayed evidence of positive selection, with one, *CSNK1A1-ENSG00000230551*, having selection within its core region, and the other two, *FLT1* and *SALL4*, showing selection within their larger introgressing segment.

We were able to identify that variants overlapping genes previously described as being related to reproduction within humans and mice show pleiotropic expressions, with genome-wide significant markers exhibiting a wide array of traits. As discussed above, other studies have documented evidence of purifying selection across the genome against archaic sequences (Fu et al., 2016; Harris & Nielsen, 2016; Juric et al., 2016) and this may explain to some extent the absence of significant GWAS results for reproductive traits for the archaic variants identified in our analysis. Nonetheless, we were able to identify 15 high-frequency archaic variants across four genes that reduce the risk of prostate cancer when considering the archaic allele. Additional evidence for archaic alleles potentially being important in cancer comes from our enrichment analysis where gene sets 1-4 all have significant enrichment for pathways previously implicated in either cancer development or suppression. We reported that adaptively introgressed variants are significantly enriched in gene pathways related to development of muscle and heart tissues and with the Notch signaling pathway, which has been previously described to be associated with proper foetal organ development. Lastly, we identified 308 archaic eQTLs that regulate gene expression levels across a variety of tissues. Over 74% of these archaic eQTLs (n=228) regulate 38 genes with expression in the ovaries, prostate, testes, uterus, and vagina for 234 genes. This may suggest that archaic haplotypes may play a currently unidentified role, or assist in connection with other processes, in developmental pathways crucial in embryonic development and later assist in other important pathways throughout a person’s lifetime.

Our analysis contributes to the study of archaic introgression and its impact on traits within the modern human genome. To our knowledge, this is the first assessment of positive selection and adaptive introgression in segments of the modern human genome known to be introgressed from archaic hominins that also overlap genes associated with reproduction. Our results describe an intriguing relationship for archaic variants within a modern human genetic background. These variants are interesting due to their strong associations with developmental related pathways, cancer incidence, and potential roles in helping indirectly balance important endogenous levels that have been previously tied to fertility outcomes. Despite these results, future work is necessary to fully understand the connections between these variants and their roles within the human genome. In conclusion, our work has clearly outlined the adaptive value of a number of variants and genes that are the direct result of admixture with archaic hominins and the roles they may have regarding human health, disease, and regulating normal development.

## Data availability statement

The SPrime software is available on GitHub [at this link](https://github.com/browning-lab/sprime) and access to the RAiSD software is also available on GitHub [here](https://github.com/alachins/raisd). The SPrime protocol, and subsequent links to relevant software used in the protocol, [are available here](https://www.sciencedirect.com/science/article/pii/S2666166721002574). The 1KGP + HGDP call set, and associated metadata tables were acquired from the [gnomAD website](https://gnomad.broadinstitute.org/downloads#v3-hgdp-1kg-tutorials). The Altai Neanderthal, Vindija Neanderthal, and Denisovan VCF files can be downloaded [directly here](http://cdna.eva.mpg.de/neandertal/Vindija/VCF/), and their associated mask files [at this link](http://cdna.eva.mpg.de/neandertal/Vindija/FilterBed/). The Chagyrskaya Neanderthal VCF file and mask files can be found [here](http://cdna.eva.mpg.de/neandertal/Chagyrskaya/). High resolution figures and supplemental tables are available at https://doi.org/10.5281/zenodo.14047487.

